# kb_DRAM: Annotating and functional profiling of genomes with DRAM in KBase

**DOI:** 10.1101/2022.03.13.484145

**Authors:** Michael Shaffer, Mikayla A Borton, Ben Bolduc, Parsa Ghadermazi, Janaka N Edirisinghe, José P Faria, Christopher S Miller, Siu Hung Joshua Chan, Matt B Sullivan, Christopher S Henry, Kelly C. Wrighton

## Abstract

**Summary:** Annotation is predicting the location of and assigning function to genes in a genome. DRAM is a tool developed to annotate bacterial, archaeal and viral genomes derived from pure cultures or metagenomes. DRAM distills multiple gene annotations to summaries of functional potential. Despite these benefits, a downside of DRAM is processing requires large computation resources, which limits its accessibility, and it did not integrate with downstream metabolic modelling tools. To alleviate these constraints, DRAM and the viral counterpart, DRAM-v, are now available and integrated in the freely accessible KBase cyberinfrastructure. With kb_DRAM users can generate DRAM annotations and functional summaries from microbial or viral genomes in a point and click interface, as well as generate genome scale metabolic models from these DRAM annotations.

**Availability and Implementation:** The kb_DRAM software is available at https://github.com/shafferm/kb_DRAM. The kb_DRAM apps on KBase can be found in the catalog at https://narrative.kbase.us/#catalog/modules/kb_DRAM. A narrative with examples of running all KBase apps is available at https://kbase.us/n/88325/84/.

**Contact:** Michael Shaffer; Kelly Wrighton

**Supplementary Information:** Supplementary data are available at Bioinformatics online.

## Introduction

Genome annotation is gene prediction and then attaching biological functional assignment to genes. Protein coding sequences are commonly assigned function via homology searches to protein databases which contain sequences with assigned or inferred functional content. A number of genome annotators targeting microbial genomes have been developed (Tanizawa *et al*., 2018; Seemann, 2014; Dong and Strous, 2019; Aziz *et al*., 2008; Zhou *et al*., 2020). We previously developed DRAM, a genome annotator which allows the user to compile annotations from multiple functionally divergent protein databases at one time, then synthesizes this content into functional profiles for each genome (Shaffer *et al*., 2020). This allows the user to rapidly understand the collection of biologically functions encoded in a set of microbial genomes.

DRAM is limited by the high computational requirements of rapidly searching against large protein databases. It requires a minimum of 128 GB of RAM to set up and 64 GB of RAM to annotate. Thus, the use of DRAM is currently limited to those with access to large compute servers. Recently cyberinfrastructure platforms have been built that provide access to computing resources as well as point-and-click interfaces to software that would usually require command line access (Merchant *et al*., 2016; Arkin *et al*., 2018; Afgan *et al*., 2018).

Genome scale metabolic models (GEMs) are representations of the metabolic reactions that occur within a bacterial cell. These reactions can be predicted from genome annotations, but many bacterial genome annotators do not generate output that is easily integrated into modeling frameworks. Additionally, recent research has shown value in combining annotations to improve genome function coverage (Griesemer *et al*., 2018), making frameworks with interoperable support of genome annotators and GEM construction more valuable.

We have built a DRAM KBase module (kb_DRAM). Here we show that using kb-DRAM, anyone with access to the KBase cyberinfrastructure (Arkin *et al*., 2018) can annotate microbial genomes with DRAM, distill these annotations into visualizations of predicted genomic functions, and use the annotations to build GEMs. Also, using bacterial genomes derived from phylogenetically distinct lineages, we show value added of including kb_DRAM annotation alongside an established KBase annotator, RAST (Aziz *et al*., 2008).

## kb_DRAM

KBase is a cyberinfrastructure platform which allows users to use common bioinformatics tools to analyze public data or a user can upload their own. Within KBase users can process microbial genomics and metagenomics data from raw reads to assemblies and bins or imported data from other sources. kb_DRAM is a plugin that provides three KBase apps. These can (1) annotate microbial DNA sequences from assemblies, isolate genomes or metagenome assembled genomes (KBase assembly objects), (2) annotate predicted coding sequences from microbial genomes (KBase genome objects) or (3) annotate viral genomes identified form metagenomes using DRAM-v.

The kb_DRAM apps use the same databases as the default DRAM installation (KOfam (Aramaki *et al*., 2020), dbCAN2 (Zhang *et al*., 2018), PFAM (El-Gebali *et al*., 2019), MERPOS (Rawlings *et al*., 2018)) to annotate predicted microbial protein coding genes as well as barrnap (https://github.com/tseemann/barrnap) for rRNA identification and tRNA-scanSE (Chan and Lowe, 2019) for tRNA detection. The DRAM product is an interactive heatmap that highlights the functional potential of genomes or metagenomes. With the microbial genome and metagenome annotation apps the product is shown in the KBase narrative so the user can understand genome function within the browser. All files generated by DRAM (e.g. raw annotations, distillate, genome completion) are available to download so users can understand their genomes more deeply. kb_DRAM apps also generate annotated KBase genome objects, which can be used in downstream analyses in KBase including building GEMs.

DRAM-v is designed to annotate viral genomes that are identified using VirSorter (Roux *et al*., 2015). DRAM-v uses the same functional databases as DRAM with the addition of RefSeq viral. To annotate with DRAM-v within KBase users can start with metagenomic assemblies and identify potential viral contigs from metagenomes using the VirSorter app in KBase. The output of the VirSorter app is then passed to the DRAM-v app for AMG annotation. The DRAM-v app shows the interactive product heatmap, which highlights potential AMGs identified in the dataset along with confidence scores for each and allows the user to download all other DRAM-v files.

## DRAM annotations of microbial genomes can generate quality GEMs in KBase

To demonstrate DRAM in KBase and show its compatibility with downstream applications, we annotated two genomes with the RAST and DRAM apps. *E. coli* was chosen as a well characterized bacterial genome, while *P. normanii*, a member of the candidate phyla radiation, represents a genome obtained from uncultivated microbes through metagenomics with limited functional curation (Castelle *et al*., 2018). Full narratives with these data are available on KBase (E. coli: https://kbase.us/n/103341/23/, P. normanii: https://kbase.us/n/95384/52/). For both genomes, when comparing the outputs of DRAM to RAST, DRAM annotations yielded more ModelSEED reactions, a measure of how many gene annotations could be converted to metabolic reactions (Fig. 1A, Supplementary table 1). As expected, due to depth of study of *E. coli*, we obtained more reaction specific annotations for *E. coli* and much fewer for *P. normanii*. Ultimately, as others have shown (Griesemer *et al*., 2018), there was value in using more than one annotator, as merged DRAM + RAST annotations yielded 1.5x and 1.4x more reactions than DRAM and 1.6x and 2.3x for RAST in *E. coli* and *P. normanii*, respectively.

**Figure 1:**
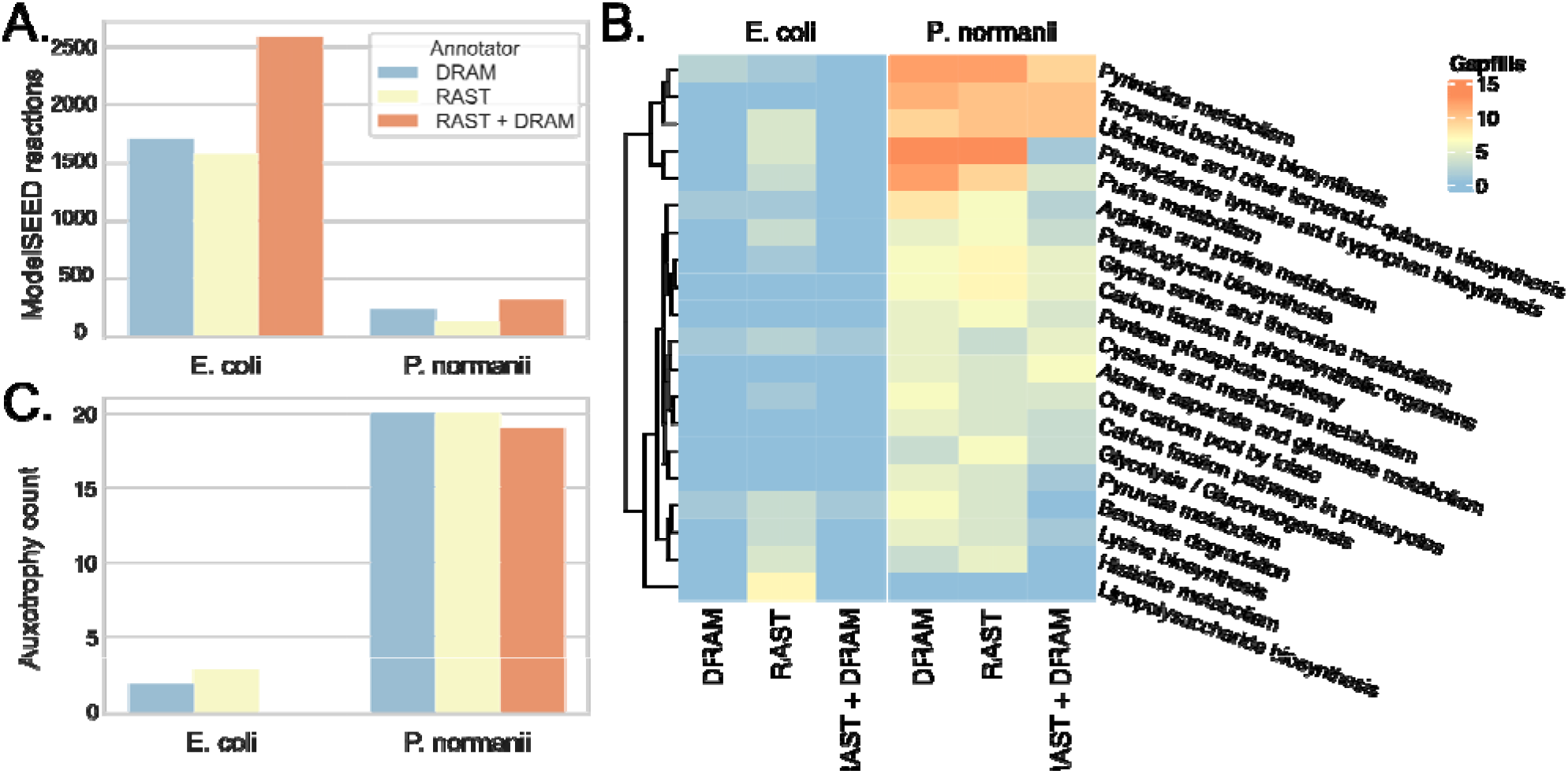
Annotation and modelling performance on three genomes. A. Number of modelSEED reactions assigned by each tool on each genome. B. Heatmap showing number of gap filled reactions required for growth on glucose minimal media per pathway. Only pathways with >=5 gap fills required by at least one annotation set shown. C. Number of auxotrophies predicted present based on annotations from each tool for each genome.

Next, we constructed GEMs using the RAST, DRAM, and DRAM + RAST annotations and GEMs were gap filled using glucose minimal media (see Supplementary Methods). In *E. coli* RAST had more reactions in the model than DRAM but fewer in *P. normanii*. In both cases DRAM + RAST outperformed each annotator alone (Supplementary table 1). We failed to find a clear pattern of better performance by DRAM or RAST in any particular metabolic pathway (Figure 1B). Subsequently, the GEMs were characterized to predict auxotrophies. In all genomes the merged annotation model showed the least number of auxotrophies (Figure 1C). Interestingly for the *E. coli* model both RAST and DRAM predicted auxotrophies, while merging the annotations removed these auxotrophies, yielding a final GEM more consistent with expected experimental evidence (Tao *et al*., 1999). We note that a large number of auxotrophies are still predicted for *P. normanii*, even when merging annotations. This finding may be biological, reflecting the symbiotic lifestyle predicted for members of this species, as many of these reactions could be provided by the host (He *et al*., 2021). However, there is no experimental data for *P. normanii* to validate this inference at this time.

## Conclusion

Here we present a KBase module with apps for running DRAM and DRAM-v. This resource enables computationally intensive genome annotation by broader audiences. We highlight that both DRAM-v and DRAM are integrated into the KBase cyberinfrastruture with the ability to ingest data from and pass data to other KBase applications. We show that DRAM can be applied to generate gene annotations from phylogenetically distinct genomes derived from pure cultures and metagenomics. Using *E. coli*, we demonstrated the addition of DRAM annotations yielded a GEM that was consistent with experimental evidence. Thus, the addition of DRAM and DRAM-v to KBase is expected to enhance user analyses of genome function and have clear value when transferring gene annotations to modeling frameworks.

## Acknowledgements

KCW was funded by an Early Career Award from the U.S. Department of Energy Office of Science, Office of Biological and Environmental Research, under award number DE-SC0019746. This work was also funded by grant DE-SC0021350 from U.S. Department of Energy Office of Science, Office of Biological and Environmental Research. We would also like to thank Elisha M Wood-Charlson.

